# Singletrack: An Algorithm for Improving Memory Consumption and Performance of Gap-Affine Sequence Alignment

**DOI:** 10.1101/2025.10.31.685625

**Authors:** Lorién López-Villellas, Cristian Iñiguez, Albert Jiménez-Blanco, Quim Aguado-Puig, Miquel Moretó, Jesús Alastruey-Benedé, Pablo Ibáñez, Santiago Marco-Sola

## Abstract

**Motivation:** Advances in DNA sequencing have outpaced advances in computation, making sequence alignment a major bottleneck in genome data analyses. Classical dynamic programming (DP) algorithms are particularly memory-intensive, especially when computing gap-affine and dual gap-affine alignments. Existing strategies to reduce memory consumption often sacrifice either speed or alignment accuracy.

**Results:** We present Singletrack, an efficient algorithm for backtrace gap-affine and dual gap-affine alignments that requires only storing a single DP matrix. Compared to classical DP algorithms, Singletrack removes the need to store additional matrices (i.e., 2 for gap-affine and 4 for dual gap-affine), significantly reducing memory consumption and, in turn, reducing pressure on the memory hierarchy and improving overall performance. Most importantly, Singletrack is a general backtrace method compatible with state-of-the-art DP-based algorithms and heuristics, such as the Suzuki-Kasahara (SK) and the Wavefront Alignment (WFA) algorithms. Our results demonstrate that Singletrack accelerates the SK implementation of KSW2, used within Minimap2, by up to 1.4×. Similarly, Singletrack enhances the performance of the WFA implementation in WFA2-lib by 1.2–2.1× while reducing memory usage by 3× for gap-affine and 5× for dual gap-affine. Compared to the efficient linear-memory BiWFA algorithm, the Singletrack-accelerated version of WFA trades a practical increase in memory usage for up to 5.2× higher performance.

**Availability:** All the implementations of the Singletrack algorithm presented in this work are available at https://github.com/LorienLV/singletrack.

## 1 Introduction

The rapid advancements in DNA sequencing technologies over the past years have enabled the generation of vast amounts of sequencing data at unprecedented speeds (Goodwin *et al*., 2016). The massive surge in sequencing data production has introduced a computational bottleneck in large-scale sequence analyses (Pfeifer, 2016; Alser et al., 2020, 2022). Notably, sequence alignment remains one of the most computationally demanding tasks in modern bioinformatics pipelines, such as read mapping (Alser *et al*., 2021; Schbath et al., 2012), de novo assembly (Liao *et al*., 2019; Jain et al., 2018), and pangenome construction (Garrison *et al*., 2024; Wang et al., 2022).

The goal of pairwise sequence alignment is to find a sequence of operations (i.e., match, mismatch, insertion, and deletion) that maps one sequence to another while minimizing a given cost function. In practice, the gap-affine and dual gap-affine cost functions are preferred due to their ability to model biologically accurate alignments.

Classical dynamic programming (DP) algorithms are commonly used to compute gap-affine and dual gap-affine alignments. In their basic formulation, these algorithms require computing and storing multiple DP matrices (i.e., 3 for gap-affine and 5 for dual gap-affine), each of size (*n* + 1) × (*m* + 1), where *n* and *m* are the input sequence lengths. These matrices are later traced back to recover the optimal alignment.

Building upon this classical formulation, several modern alignment algorithms have been proposed to improve execution time and reduce memory usage. One notable example is the Suzuki-Kasahara (SK) formulation(Suzuki and Kasahara, 2018), which reformulates the gap-affine scoring equations so that the matrices store score differences instead of absolute scores. This formulation increases the number of matrices to store, but reduces the size of each cell, resulting in overall memory savings. This approach is used by KSW2(Li, 2018a), at the core of Minimap2 (Li, 2018b), and by the libgaba (Suzuki and Kasahara, 2018) library. More recently, the Wavefront Alignment Algorithm (WFA) (Marco-Sola *et al*., 2020) was proposed, which exploits homologous regions between sequences to substantially reduce the number of DP matrix cells computed and lower the memory requirements, particularly when aligning highly similar sequences.

Nevertheless, these implementations store all the computed cells of the DP matrices. In both the classical formulation and WFA, this involves storing three matrices for gap-affine and five for dual gap-affine alignment. In the case of the SK formulation, it requires storing four matrices for gap-affine alignment and six for dual gap-affine alignment. Compared to simpler cost functions, such as edit distance, using multiple matrices significantly increases memory requirements, making these algorithms particularly memory-intensive and degrading overall performance, especially when scaling to long sequences or when processing large alignment batches.

To address this challenge, practical sequence alignment implementations for gap-affine and dual gap-affine scoring adopt a variety of techniques to reduce memory usage. These include using heuristics at the expense of losing accuracy, storing compact backtrace representations instead of full matrices, and space-time trade-offs, such as divide-and-conquer strategies that recompute partial results to avoid storing the DP matrices entirely.

Heuristic-based pruning is a common technique to reduce memory usage by restricting the number of DP cells computed and stored. For instance, static banding restricts DP computations to a fixed diagonal band around the main diagonal, where the optimal alignment is expected to be found (Ukkonen, 1985). Alternatively, the computation region can be adjusted dynamically during alignment, pruning cells that fall too far from the most promising paths (Altschul, 1997; Li, 2016). Such heuristics may drastically reduce memory usage but risk missing the optimal alignment if it lies outside the explored region. Notwithstanding, they are standard (usually optional) features in many libraries, including KSW2 (Li, 2018a), Edlib (Šošić and Šikić, 2017), Parasail (Daily, 2016), and WFA2-lib (Marco-Sola *et al*., 2020).

Another memory-saving technique involves storing a backtrace matrix instead of the full DP matrices during alignment. That is, the aligner keeps only those DP cells required to compute subsequent DP cells, while the backtrace information (i.e., the direction of the next DP cell during the backtrace) is stored in a compact auxiliary matrix using small-width elements (e.g., 8-bit values). This approach is implemented in widely-used libraries such as KSW2 (Li, 2018a) and Parasail (Daily, 2016). While effective at reducing memory consumption, this method introduces significant overhead from additional data manipulation and memory accesses during the alignment process.

Another common approach is to trade additional computation for reduced memory usage using a divide-and-conquer approach. The divide-and-conquer paradigm can be leveraged to recursively break the alignment process into smaller pieces, effectively transforming the quadratic memory consumption of classical algorithms to linear memory consumption. This technique is used by Hirschberg’s algorithm (Hirschberg, 1975) and Bi-WFA (Marco-Sola *et al*., 2023), a version of WFA capable of exploiting divide-and-conquer. However, these linear-memory approaches, while extremely memory-efficient, tend to be slower than their quadratic-memory counterparts for moderate sequence lengths. For instance, the BiWFA paper reports that WFA outperforms BiWFA for sequences of up to 100 Kbp.

This work presents Singletrack, an efficient backtracing algorithm that requires storing only a single DP matrix, even when computing gap-affine and dual gap-affine alignments. We demonstrate that storing a single DP matrix is enough to recover the optimal alignment while maintaining the linear time complexity of the backtrace. Moreover, we show that Singletrack is broadly applicable with state-of-the-art DP-based methods, such as SK formulation, WFA, and commonly used heuristic strategies. To demonstrate applicability and performance, we integrated Singletrack into KSW2 (which implements SK) and WFA2-lib (which implements WFA), yielding the KSW2+Singletrack and WFA+Singletrack optimized implementations. As a result, we demonstrate that Singletrack enables equal memory consumption for gap-affine and dual gap-affine alignments in KSW2, while reducing memory usage by 3× and 5× for gap-affine and dual gap-affine, respectively, in WFA2-lib. Beyond memory savings, Singletrack also improves performance, delivering speedups of 1.2× to 2.1× compared to the baseline implementations. Compared to the linear-space BiWFA algorithm, WFA+Singletrack achieves speedups of up to 5.2× at the cost of increased memory usage, making it a compelling alternative for moderately long sequences.

## 2 Background

### 2.1 Pairwise Sequence Alignment

Let the query *q* = *q*_0_, *q*_1_, …, *q*_*n*−1_ and the target *t* = *t*_0_, *t*_1_, …, *t*_*m*−1_ be two sequences of length *n* and *m* characters, respectively. The optimal sequence alignment is the sequence of four basic operations (i.e., match, mismatch, insertion, and deletion) that transform *q* into *t* while minimizing a given cost function.

Naturally, the choice of cost function strongly affects the alignment result and its biological relevance, as well as the computational and memory requirements. The simplest cost functions are the edit-distance and gap-linear (a.k.a. weighted edit-distance). These functions use three values to score matches (*a*), mismatches (*x*), and gap extensions (*e*) for both insertions and deletions. In the case of the edit-distance, these values are fixed to *a* = 0, *x* = 1, *e* = 1, while gap-linear functions allow customizable values to match biological or empirical scoring needs. In the following, for simplicity, we define a substitution function *S*(*i, j*) that evaluates to *a* if *q*_*i*−1_ = *t*_*j*−1_ and *x* if *q*_*i*−1_ ≠*t*_*j*−1_.

Edit-distance and gap-linear alignments are traditionally computed using DP algorithms that require computing and storing a single DP matrix *M* of size (*n* + 1) · (*m* + 1), where each cell *M*_*i,j*_ is filled using a simple recurrence, *M*_*i,j*_ = min{*M*_*i,j*−1_ + *e, M*_*i*−1,*j*_ + *e, M*_*i*−1,*j*−1_ + *S*(*i, j*)}. As a result, the optimal score of the alignment is found in *M*_*n,m*_, and the optimal alignment can be recovered by tracing back from this cell the sequence of minimum-cost operations that led to it, using the backtrace algorithm.

### 2.2 Gap-Affine and Dual Gap-Affine Alignment

Despite their simplicity, gap-linear cost functions have notable limitations. These functions model long gaps as multiple independent insertions or deletions, whereas in reality, they often stem from a single biological event. This can lead to fragmented alignments and misleading interpretations, making linear models unsuitable in some cases. To reflect this phenomenon, the gap-affine function (Smith *et al*., 1981; Gotoh, 1982) introduces a gap-opening cost (*o*_1_). This function models cases where a gap follows a different type of operation, such as a mismatch or match. For example, two insertions separated by a non-insertion costs 2 · *o*_1_ + 2 · *e*_1_, while a single gap of two insertions costs *o*_1_ + 2 · *e*_1_.

However, gaps in sequence alignments arise from two distinct mechanisms: evolutionary events, which can introduce long gaps through a single event (e.g., transposon insertion), and sequencing errors, which tend to generate multiple short gaps. To model this distinction and produce more biologically accurate alignments, the dual gap-affine function (Gotoh, 1990) is often preferred. To that end, the dual gap-affine function introduces a second set of gap penalties (*o*_2_, *e*_2_), allowing for two types of gaps. One type of gap has a low opening cost but a high extension cost (to model short gaps), while the other has a high opening cost but a low extension cost (to model large gaps). For each gap of length *l*, the alignment algorithm chooses the lower-cost option: min{*o*_1_ + *l* · *e*_1_, *o*_2_ + *l* · *e*_2_}.

Similar to gap-linear models, gap-affine and dual gap-affine alignments are typically computed using traditional DP algorithms. However, computing gap-affine alignments requires computing and storing three DP matrices (*M*,*I*1, and *D*1). More demanding still, dual gap-affine alignment requires five matrices (*M*,*I*1,*D*1,*I*2, and *D*2). In particular, the *I* matrices (*I*_1_ and *I*2) and *D* matrices (*D*_1_ and *D*2) track optimal scores for alignments ending in insertions and deletions, respectively. Then, the *M* matrix summarizes the minimum alignment score between prefixes *q*_0,…,*i*−1_ and *t*_0,…,*j*−1_ for each cell *M*_*i,j*_. The recurrence equation used to compute the gap-affine DP matrices is presented in Eq. 1. Additional details, including initial conditions and the extension to the dual gap-affine cost function, are provided in the Supplementary Material.

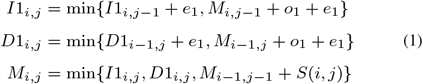

The optimal score of the alignment is found in *M*_*n,m*_. The subsequent backtrace process utilizes the information in all the matrices to retrieve the path that led to *M*_*n,m*_.

As in gap-linear models, the optimal alignment score is located in cell *M*_*n,m*_. To reconstruct the alignment, a backtrace procedure traverses the *M, I*, and *D* matrices, following the path of minimum-cost operations that led to this cell (i.e., optimal alignment).

### 2.3 Classical Backtrace Algorithm

The classical backtrace algorithm reconstructs an optimal alignment path by traversing the *M, I*, and *D* matrices. More in detail, the backtrace procedure starts at the last cell, *M*_*n,m*_. At each step, the algorithm identifies the predecessor cell that originated the score of the current cell by evaluating the transitions defined in Eq. 1. This transition corresponds to an alignment operation, belonging to the optimal alignment, which is commonly represented using the CIGAR string format: `M` for match, `X` for mismatch, `I` for insertion, and `D` for deletion. The algorithm then moves to the identified predecessor cell and repeats the process until it reaches the starting cell *M*_0,0_. It is important to note that multiple transitions may lead to the same minimum-cost cell, resulting in different equally optimal alignment paths. In the case of the dual gap-affine function, the backtrace algorithm is analogous.

For example, Figure 1 shows how the classical backtrace algorithm operates over the computed DP matrices when aligning two sequences (*q* = `GCA` and *t* = `GCCAA`) using a gap-affine cost function with penalties *a* = 0, *x* = 4, *o*_1_ = 6, and *e*_1_ = 2. The dark-shaded cells in the figure indicate the optimal alignment path. The backtrace begins at cell *M* [*n, m*] = *M* [3, 5] = 10. The only valid transition is from *M* [2, 4], since *M* [3, 5] = *M* [2, 4]+*S*(3, 5). The current cell is updated to *M* [2, 4], and the CIGAR string is set to `M`. From *M* [2, 4], the algorithm moves to *I*_1_[2, 3], as *M* [2, 4] = *I*_1_[2, 3] + *e*_1_, updating the CIGAR string to `IM`. Then, since *I*_1_[2, 3] = *M* [2, 2] + *o*_1_ + *e*_1_, the path returns to *M* [2, 2], resulting in `IIM`. Next, *M* [2, 2] traces back to *M* [1, 1] via *M* [2, 2] = *M* [1, 1] + *S*(2, 2), giving `MIIM`. Finally, *M* [1, 1] = *M* [0, 0] + *S*(1, 1), and the completed CIGAR string is `MMIIM`. This example illustrates how the presence of a gap in the alignment requires transitioning into *I*_1_ or *D*_1_ during backtrace and returning to *M* once the gap is closed.

**Figure 1.**
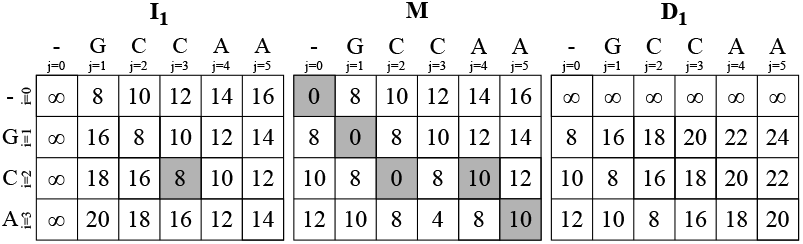
Classical backtrace for the gap-affine alignment of *q* = `GCA` and *t* = `GCCAA` using penalties *a* = 0, *x* = 4, *o*_1_ = 6, and *e*_1_ = 2. Dark-shaded cells indicate the optimal path over matrices *M, I*_1_, and *D*_1_.

## 3 The Singletrack Algorithm

We now describe Singletrack, a method for performing the backtrace of gap-affine and dual gap-affine alignments in linear execution time using only the *M* matrix. As a result, the alignment process no longer needs to store the indel matrices, significantly reducing memory usage. Without loss of generality, we focus on the gap-affine case for simplicity, noting that the method extends naturally to dual gap-affine alignments.

We observe that Equation 1 indicates that while computing *M* requires values from all three matrices, each indel matrix (*I*_1_, *D*_1_) depends only on *M* and itself. Thus, any cell in an indel matrix traces back either to a cell in *M* or to another cell within the same indel matrix. Hence, during backtrace, if the current cell is in an indel matrix such as *I*_1_, we can avoid accessing *I*_1_ by checking first whether *s* = *M*_*i,j*−1_ − *o*_1_ − *e*_1_. If this holds, the gap was opened from *M*_*i,j*−1_; otherwise, the gap continues, and the predecessor is necessarily *I*1_*i,j*−1_ with score *s*^*′*^ = *s* − *e*_1_. As a result, backtracing through *I*_1_ can be performed iteratively without accessing its scores, until the trace returns to a cell in *M*.

Singletrack generalizes this idea to avoid accessing *I*1 and *D*1 scores. When the current cell is in *M*, say *M*_*i,j*_, we first check if the predecessor is in *M* by checking if *M*_*i,j*_ = *M*_*i*−1,*j*−1_ + *S*(*i, j*). If not, the predecessor cell is either *I*1_*i,j*−1_ or *D*1_*i*−1,*j*_. Although we do not know which one, from Equation 1, we know that the score of the predecessor cell must be *s*^*′*^ = *M*_*i,j*_ − *e*_1_. At this point, we explore two tentative backtrace paths simultaneously: one assuming the predecessor lies in *I*_1_, and the other in *D*_1_. Then, we extend both paths step by step, increasing the gap length *l* and computing the expected score to return to *M* as *s*(*l*) = *M*_*i,j*_ − *o*_1_ − *l* · *e*_1_. Eventually, either *M*_*i,j*−*l*_ = *s*(*l*) or *M*_*i*−*l,j*_ = *s*(*l*) holds, indicating that the correct backtrace path goes from *M*_*i,j*_ to *M*_*i,j*−*l*_ through *I*_1_ (insertion) or to *M*_*i*−*l,j*_ through *D*_1_ (deletion), respectively. Once a consistent path back to *M* is found, the other path is discarded, and the algorithm continues the backtrace in *M*. The pseudocode of Singletrack’s backtrace algorithm is shown in Algorithm 1.

Following the previous example, Figure 2 shows the behaviour of the Singletrack backtrace algorithm. For that, we show both the stored and recomputed cells: scores stored in memory (corresponding to the *M* matrix) are displayed in black, while tentative scores, computed on the fly during the backtrace but not stored, are shown in dark grey. In the first step, the behaviour matches the classical algorithm since *M*_3,5_ = *M*_2,4_ + *S*(3, 5) and the CIGAR string is set to `M`. At *M*_2,4_, since *M*_2,4_*≠M*_1,3_ + *S*(2, 4), the predecessor cell must be either *I*1_2,4_ or *D*1_2,4_, both having a tentative score *s*^*′*^ = *M*_2,4_ = 10. For a tentative gap length of *l* = 1, the expected score back to *M* should be *s*^*′*^ = *M*_2,4_ − *o*_1_ − *l* · *e*_1_ = 2, which does not correspond to *M*_2,3_ or *M*_1,4_. Note that the tentative score of *I*1_2,3_ and *D*1_1,4_ is *s*^*′*^ = *M*_2,4_ − *e*_1_ = 8, corresponding to the true score at *I*1_2,3_ (correct backtrace path) but not of *D*1_1,4_ (see Figure 1). Increasing the gap length to *l* = 2, the algorithm extends both paths and checks whether the updated score back to *M, s*^*′*^ = 10 − 6 − 2 · 2 = 0, matches either *M*_2,2_ or *M*_1,3_. Since *s*^*′*^ = *M*_2,2_, the correct path goes through *I*1, and two insertions are added to the CIGAR string, which becomes `IIM`. From this point, the algorithm proceeds as in the classical case, and the final CIGAR string is `MMIIM`, matching the result produced by the original backtrace algorithm.

**Figure 2.**
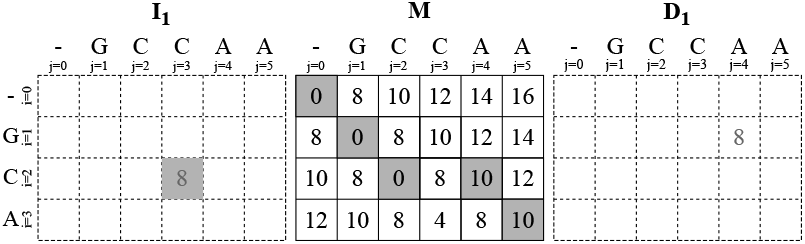
Singletrack backtrace for the gap-affine alignment of *q* = `GCA` and *t* = `GCCAA` using penalties *a* = 0, *x* = 4, *o*_1_ = 6, and *e*_1_ = 2. Only the matrix *M* is stored. Strictly required *I*_1_ and *D*_1_ values are recomputed on the fly. Dark-shaded cells indicate the optimal path.

### Algorithm 1

Singletrack gap-affine backtrace

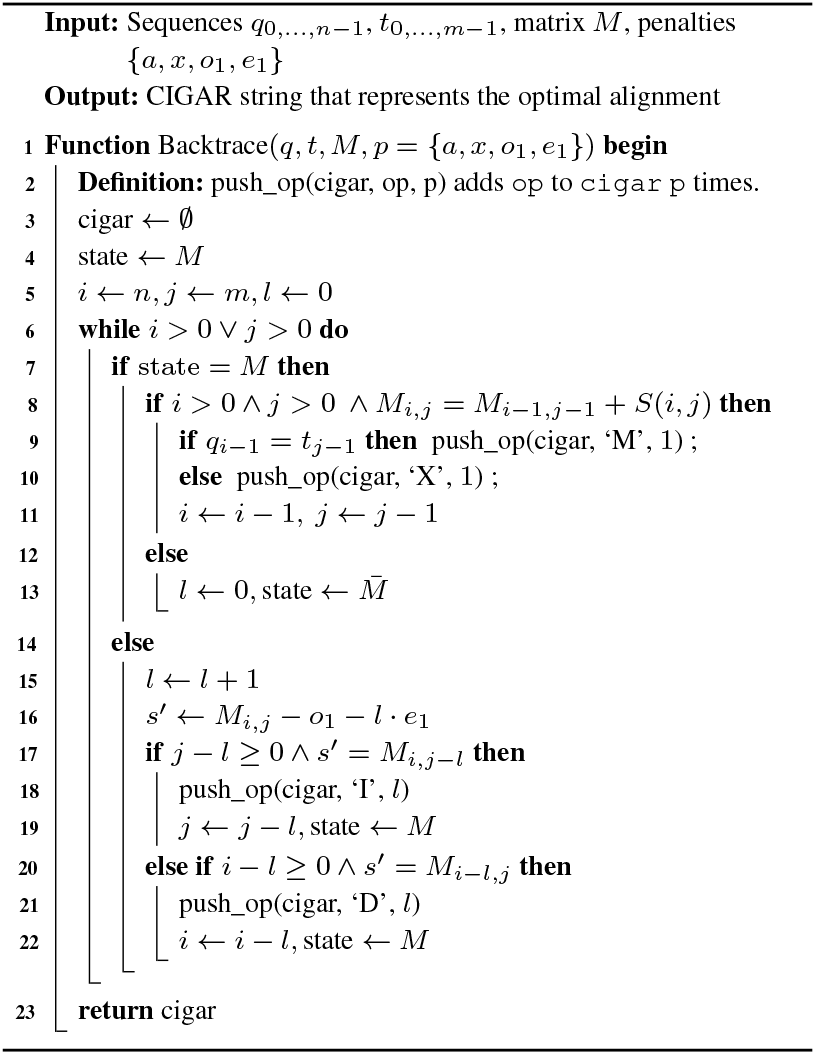

The time complexity of the Singletrack algorithm remains linear, *O* (*n*+ *m*), although it may explore up to twice as many cells as the classical algorithm in the worst case. Nevertheless, the backtrace step generally constitutes a small fraction of the total alignment time. In the case of dual gap-affine alignments, the approach is identical, but with four tentative paths corresponding to *I*1, *D*1, *I*2, and *D*2, potentially increasing the worst-case number of explored cells by up to four times compared to the classical algorithm.

The key advantage of Singletrack lies in its ability to compute optimal alignments while storing only the *M* matrix and a minimal set of values from the other matrices (i.e., *I*1, *D*1, *I*2, *D*2). In particular, to compute *M* values using Equation 1 column-wise, the algorithm only requires storing a row from *I*_1_ (and *I*2) and a single cell from *D*_1_ (and *D*2). Ultimately, this results in a memory reduction of approximately 3× for gap-affine and 5× for dual gap-affine alignments, without adding any overhead to the initial alignment computation.

Moreover, requiring only a small set of the indel matrices enables software optimizations. For instance, the single element needed from *D*1 (and *D*2) can be stored directly in a CPU register rather than in memory, significantly reducing memory accesses and latency, and improving performance. Similarly, the memory of the single column required from *I*1 (and *I*2) can be reused continuously, which improves the cache locality and reduce pressure on lower levels of the memory hierarchy, which is particularly advantageous when aligning long sequences or processing large alignment batches.

Furthermore, Singletrack is fully compatible with all the heuristic techniques commonly used in sequence alignment. Strategies such as static or adaptive banding, early termination, drops, or score filtering can be integrated into the alignment and backtrace processes without affecting correctness.

More importantly, Singletrack can be applied to any DP-based sequence alignment algorithm, provided that the required scores of the *M* matrix are available during the backtrace. As shown in the following sections, Single-track integrates seamlessly with the Suzuki-Kasahara formulation (Suzuki and Kasahara, 2018) and with the Wavefront Alignment Algorithm (Marco-Sola *et al*., 2020). Additionally, it is compatible with heuristic alignment approaches, such as bands, requiring only additional checks for boundary conditions during the backtrace.

We have integrated the Singletrack backtracking into the classical dynamic programming algorithm. The implementation is available at https://github.com/LorienLV/singletrack.

## 4 Singletrack in Suzuki-Kasahara Formulation

The Suzuki-Kasahara (SK) formulation (Suzuki and Kasahara, 2018) extends the difference encoding idea from the BPM algorithm (Myers, 1999) to the gap-affine cost function, storing only the differences between adjacent DP cells rather than full scores. Since these differences and alignment penalties are usually small integers, they can be efficiently represented with reduced data types, such as 8-bit integers. The standard recurrence relations (Equation 1) are reformulated in terms of score differences for the *M, I*1, and *D*1 matrices. Specifically, we define the horizontal, vertical, and diagonal differences in the *M* matrix as Δ*H*_*i,j*_ = *M*_*i,j*_ − *M*_*i,j*−1_, Δ*V*_*i,j*_ = *M*_*i,j*_ − *M*_*i*−1,*j*_, and *A*_*i,j*_ = *M*_*i,j*_ − *M*_*i*−1,*j*−1_, respectively. Additionally, the differences between the *I*1 and *D*1 matrices and the *M* matrix are given by Δ*E*1_*i,j*_ = *I*1_*i,j*_ − *M*_*i,j*_ and Δ*F* 1_*i,j*_ = *D*1_*i,j*_ − *M*_*i,j*_. The resulting Suzuki-Kasahara recurrence relations are shown in Equation 2 (see Supplementary Material for the dual gap-affine equations).

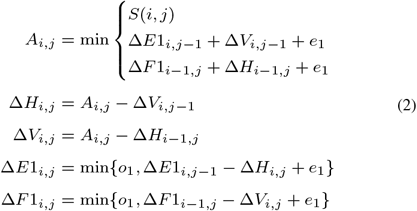

With these new recurrence relations, four matrices are stored: Δ*H*_*i,j*_, Δ*V*_*i,j*_, Δ*E*1_*i,j*_, and Δ*F* 1_*i*−1,*j*_, compared to only three in the original formulation (where *A*_*i,j*_ is a temporary variable and not stored in memory). However, since the new matrices store narrow-integer values (e.g., 8-bit integers), the overall memory footprint is actually reduced compared to the original approach (assuming 32-bit integer encoding baseline). Furthermore, this reformulation increases the number of elements that can be processed simultaneously through SIMD (Single Instruction, Multiple Data) operations available in modern CPUs, enhancing computational performance.

To apply the Singletrack backtrace under the SK formulation, it is sufficient to retain only the Δ*V* and Δ*H* matrices, as they provide all the information required to reconstruct the original *M* matrix (the remaining Δ*E*_1_ and Δ*F*_1_ matrices can be discarded). However, the Singletrack algorithm requires integer values from the *M* matrix to trace back the optimal alignment from *M*_*n,m*_ to *M*_0,0_. To address this, we define a new matrix *M* ^*′*^ such that 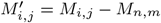 for all *i* ∈ {0, …, *n*} and *j* ∈ {0, …, *m*}. By construction, 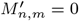, providing a known integer value to start the backtrace. During the backtrace, the relevant entries of *M* ^*′*^ are reconstructed on the fly using the stored differential matrices Δ*V* and Δ*H* as 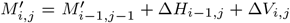 Since *M* ^*′*^ is simply a shifted version of *M*, the original alignment path is preserved, and the general Singletrack algorithm remains unchanged. These ideas naturally extend to the SK formulation of the dual gap-affine cost function.

We have integrated Singletrack into KSW2^1^, which provides a SIMD-accelerated implementation of the Suzuki-Kasahara formulation and serves as a core component of Minimap2 (Li, 2018b), a widely adopted read mapper and a foundational tool in many bioinformatics pipelines. KSW2 does not store the full DP matrices during the alignment. Instead, it employs the backtrace matrix approach previously described, in which the backtrace is performed on the fly during the alignment, and the backtrace information for each DP cell is stored in a single compressed matrix of 8-bit elements.

We produced a minimal version of KSW2 that utilizes Singletrack for backtrace, referred to as KSW2+Singletrack. Instead of computing the backtrace matrix, the matrices Δ*H* and Δ*V* are stored. This approach requires approximately twice the memory compared to the original KSW2; however, it eliminates all data manipulation associated with the backtrace matrix. Note that KSW2+Singletrack uses 2× and 4× less memory than the classical approach that stores all matrices for gap-affine and dual gap-affine, respectively. The KSW2+Singletrack implementation is available at https://github.com/LorienLV/singletrack.

## 5 Singletrack in Wavefront Alignment Algorithm

The Wavefront Alignment Algorithm (Marco-Sola *et al*., 2020) (WFA) proposes the alternative recurrence relations to compute the optimal gap-affine alignment. For each matrix (*M, I*1, *D*1) and diagonal *k* of the DP matrix, WFA computes and stores the offset (column) of the most advanced DP cell with score *z*, known as the furthest reaching point (f.r.p.). The vector of offsets for a given score *z* is called the *z*-wavefront. In the gap-affine case, WFA computes three wavefronts (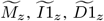) per each score value following Equation 3.

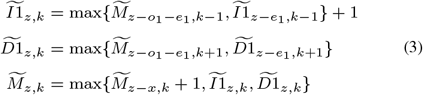

Once the wavefronts for the optimal score *s* have been computed, the alignment path is recovered through a backtrace procedure that follows the same principles as the classical backtrace algorithm. That is, the WFA backtrace reconstructs the optimal alignment path following the origin of each furthest-reaching point in Equation 3 from 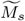 to 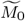, accessing all computed wavefronts in between (i.e., 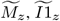 and 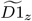)

Unlike classical DP-based alignment algorithms, which require quadratic time and memory, WFA has a time complexity of *O* (*ns*) and a space complexity of *O* (*s*^2^), where *n* is the length of the sequences (assuming *n* ∼ *m*) and *s* is the optimal alignment score. In practice, WFA outperforms most sequence alignment algorithms, especially when the sequences aligned are highly similar.

As with the other methods, the Singletrack backtrace algorithm can be applied to WFA, requiring only to store the 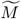 wavefronts. This process is analogous to that used in the other cases. In essence, the Singletrack algorithm follows the transitions in 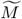 when the predecessor of the current cell is in 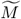, and opens tentative paths in 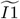 and 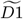 when the predecessor is not in 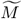 (see an extended description in the Supplementary Material). Consequently, it is not necessary to store the 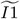 and 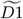 wavefronts. As in the previous sections, these ideas naturally extend to the dual gap-affine WFA.

We have integrated Singletrack into WFA2-lib^2^, which implements the Wavefront Alignment Algorithm and is utilized in several state-of-the-art genome analysis tools, including Anchorwave (Song *et al*., 2021) and wfmash (Garrison *et al*., 2024). WFA2-lib explicitly stores all the wavefronts to perform the backtrace (i.e., for each score *z*, three wavefronts in the case of gap-affine alignment and five wavefronts in the case of dual gap-affine alignment).

In our version of WFA2-lib, WFA+Singletrack, only the 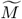 wavefronts are stored and used for the backtrace. For the indel matrices, we allocate a minimal scope of *N* wavefronts at the start of the alignment, each wavefront with the maximum possible size that may be required during the alignment. These *N* wavefronts are initially assigned to scores *z* = 0, …, *N* − 1. As the alignment progresses and a wavefront’s score moves out of scope, its memory is recycled for higher scores; i.e., the wavefronts within the scope are cycled, ensuring that no additional memory is required for the indel matrices. For sufficiently large sequences, WFA+Singletrack reduces memory usage by a factor of 3× for gap-affine and 5× for dual gap-affine alignments compared to the original WFA implementation in WFA2-lib. The modified version of WFA2-lib supporting Singletrack is available at https://github.com/LorienLV/singletrack.

## 6 Results

### 6.1 Experimental Setup

We evaluate the performance and memory consumption of KSW2+Singletrack and WFA+Singletrack, introduced in previous sections, and compare them with their original implementations. Additionally, we evaluate the divide-and-conquer version of WFA (BiWFA) as a state-of-the-art reference alignment algorithm with linear memory complexity. The classical dynamic programming implementation (with and without Singletrack) is omitted from the results due to its substantially higher computational cost compared to the optimized KSW2 and WFA algorithms.

For the evaluation, we configure all implementations to compute global alignments. When computing gap-affine alignments, penalty values are set to *a* = 0, *x* = 4, *o*_1_ = 6, and *e*_1_ = 2. For dual gap-affine alignments, penalties are set to *a* = 0, *x* = 4, *o*_1_ = 6, *e*_1_ = 2, *o*_2_ = 24, and *e*_2_ = 1.

We evaluate our method on three datasets obtained from real sequencing experiments. The first, Illumina WGS, was obtained from NIST’s Genome in a Bottle (GIAB) project^3^ and comprises 100 million sequence pairs with lengths ranging from 140 bp to 280 bp and an average error rate of 1%. The second, PacBio HiFi, was obtained from the Precision FDA Truth Challenge V2^4^ and contains 10 million sequence pairs ranging from 0.20 Kbp to 24.64 Kbp with an average error rate of 0.6%. The last dataset, ONT PromethION, was obtained from the Human Pangenome Reference Consortium (Miga and Wang, 2021) and comprises 1,312 sequence pairs with lengths of up to 309.28 Kbp. For our experiments, we select only the pairs up to 50 Kbp in length, resulting in 18 sequences with an average error rate of 12%. While the full ONT dataset was preliminarily tested, only BiWFA could complete the alignments, as the other algorithms exceeded the compute node’s memory capacity. Consequently, our evaluation focuses on the subset of sequence pairs that can be aligned by all methods under comparison, ensuring a fair and informative comparison. It is important to note that aligning such long sequences (i.e., *>* 50 Kbp) is relatively uncommon in practice, as most mappers, such as Minimap2 (Li, 2018b), typically use a seed-and-extend approach that requires aligning much smaller sequence chunks (e.g., anchors or seeds) rather than full-length sequences.

All the experiments were conducted on a computing node equipped with two Intel Xeon Platinum 8480+ processors, each featuring 56 cores and 112 threads, and 256 GiB of main memory. The main specifications of the node are summarised in Table 1.

**Table 1.** Main features of the computing nodes used to execute the experiments.

|  |  |
| --- | --- |
| Processor | 2 × Intel Xeon Platinum 8480+ |
| Cores | 2 × 56 |
| Frequency | 2 GHz |
| L1 cache (I + D) | 32 KiB + 48 KiB (per core) |
| L2 cache | 2 MiB (per core) |
| LLC | 2 × 105 MiB (shared) |
| Main Memory | 256 GiB DDR5 (16 × 16 GiB @ 4800 MHz) |

### 6.2 Single-Threaded Results

In this section, we evaluate the single-threaded performance of the original KSW2, WFA, and BiWFA implementations, compared with the Singletrack-improved variants introduced in this manuscript, KSW2+Singletrack and WFA+Singletrack, in terms of time and memory usage. Performance is evaluated by measuring the total execution time required to align complete datasets (including the backtrace phase), while memory consumption is measured as the peak resident set size (RSS) recorded during the alignment process using the GNU `/usr/bin/time` command. Table 2 summarizes the execution times and memory footprints for all five implementations using both gap-affine (aff) and dual gap-affine (2aff) cost functions. To enable comparison across datasets of varying sizes, the alignment times are normalized by the number of base pairs. For each dataset and cost function, the best-performing algorithm for each metric (performance or memory) is highlighted in bold.

**Table 2.** Single-threaded runtime and memory usage of KSW2, KSW2+ST, WFA, BiWFA, and WFA+Singletrack (aff = gap-affine; 2aff = dual gap-affine; ST = Singletrack)

| | | Time (μs) / Kbp | | | Mem (MiB) | | |
| --- | --- | --- | --- | --- | --- | --- | --- |
|  |  | Illumina | PacBio HF | PromethION | Illumina | PacBio HF | PromethION |
| aff | KSW2 | 65.8 | 2,980.6 | 7,849.8 | 3.8 | 10,463.6 | 4,534.5 |
|  | KSW2+ST | 60.8 | 3,634.5 | 10,063.8 | 4.0 | 21,864.9 | 9,065.2 |
|  | WFA | 4.8 | 13.3 | 7,447.2 | 5.3 | 427.9 | 6,337.0 |
|  | BiWFA | 8.9 | 18.6 | **4,428.1** | **3.2** | **15.1** | **17.3** |
|  | WFA+ST | **3.4** | **10.9** | 5,233.2 | 26.0 | 179.5 | 2,123.8 |
| 2aff | KSW2 | 110.5 | 4,887.9 | 12,479.2 | 3.8 | 10,463.6 | 4,537.9 |
|  | KSW2+ST | 77.7 | 4,108.7 | **11,472.8** | 4.0 | 21,864.9 | 9,066.2 |
|  | WFA | 11.3 | 50.9 | 44,884.7 | 6.8 | 2,557.0 | 37,653.1 |
|  | BiWFA | 28.5 | 172.8 | 35,223.5 | **6.0** | **58.8** | **115.4** |
|  | WFA+ST | **5.4** | **39.7** | 28,782.6 | 62.4 | 543.8 | 7,546.1 |

KSW2+Singletrack requires twice the memory of the original KSW2 for both gap-affine and dual gap-affine implementations, as it stores the Δ*H* and Δ*V* matrices instead of the backtracking matrix used in KSW. However, KSW2+Singletrack eliminates all computations and memory accesses related to constructing this backtracking matrix. This change leads to performance improvements on the Illumina dataset when using the gap-affine cost function. When aligning short sequences, both implementations have a similar memory footprint, as the total memory usage is dominated by fixed overheads such as auxiliary structures and allocation thresholds. In contrast, aligning the PacBio HF and Promethion datasets results in a slowdown of 1.2× on average due to increased memory pressure, highlighting the critical importance of memory efficiency when scaling to long sequences. However, when computing dual gap-affine (2aff), which is the primary alignment mode in Minimap2 (Li, 2018b), the memory requirements for KSW2+Singletrack remain the same as in the case of gap-affine (aff). For the baseline KSW2, constructing the backtracking matrix requires more instructions due to the additional matrices involved in dual gap-affine. As a result, KSW2+Singletrack achieves consistent speedups on all three datasets, ranging from 1.1× to 1.4×, while presenting a simpler implementation and minimal integration overhead.

Regarding WFA+Singletrack, it outperforms the original WFA, achieving 1.2–2.1× speedups and reaching the theoretical 3× and 5× memory reductions for gap-affine (aff) and dual gap-affine (2aff), respectively, on the dataset with the longest sequences (PromethION). It should be emphasized that WFA+ST does not use the same amount of memory in gap-affine and dual gap-affine, since in the latter it explores a larger number of cells in the dynamic programming (DP) matrix, i.e., the *M* wavefronts require more memory.

In terms of execution time improvement, note that WFA+Singletrack repeatedly reuses the memory allocated to the indel matrices, reducing cache misses compared to WFA, especially for larger sequences. By examining the memory usage reported in the tables, it is clear that the alignment data exceeds the capacity of both the L1D cache (48 KiB) and the L2 cache (2 MiB) across all datasets and algorithm variants. Therefore, reducing cache misses has a significant positive impact on performance.

In terms of memory consumption, WFA+Singletrack uses less memory than WFA, except for the Illumina dataset. The difference in memory usage increases as the sequence length grows, eventually reaching theoretical maximum reductions of 3× and 5×. For Illumina reads, WFA+Singletrack over-allocates memory for indel matrices assuming worst-case usage, while most of it remains unused. Despite this, it outperforms WFA 1.3× for gap-affine and 2.1× for dual gap-affine, justifying the trade-off given the low absolute memory cost.

The linear-memory BiWFA algorithm also benefits from efficient memory reuse, leading to fewer cache misses than the original WFA. However, BiWFA’s divide-and-conquer approach introduces additional overhead by recursively recomputing already-processed alignment regions. This overhead translates to WFA+Singletrack outperforming BiWFA on the Illumina and PacBio HiFi datasets for gap-affine and in all three datasets for dual gap-affine. Across the reported results, WFA+Singletrack is up to 5.2× faster than BiWFA. Although BiWFA consistently exhibits the lowest memory consumption across all datasets, the substantial performance improvements offered by WFA+Singletrack make it a compelling choice for moderately long sequences, where memory usage remains manageable.

Overall, WFA+Singletrack achieves the best performance among the evaluated algorithms and cost functions, except for the PromethION dataset, where BiWFA is faster for gap-affine and KSW2+Singletrack is faster for dual gap-affine. As expected, BiWFA consistently uses the least memory, but the higher memory consumption of KSW2+Singletrack and WFA+Singletrack is justified by their performance improvements on short to moderately large sequences.

### 6.3 Multi-Threaded Results

Modern sequence alignment tools are frequently deployed in multi-threaded settings to take advantage of today’s multi-core architectures. This is especially true in large-scale applications such as read mapping. In these contexts, scalability across multi-core computing nodes becomes a key performance metric. In this section, we evaluate the parallel performance of KSW2, KSW2+Singletrack, WFA, BiWFA, and WFA+Singletrack when executed across multiple cores on our computing node.

In this comparison, each core independently aligns the entire dataset without synchronization, thereby eliminating the effects of load imbalance or inter-thread communication on the measured performance. Execution time is calculated as the average runtime across all threads and normalized by dividing by the number of threads, yielding the scaled time. To maintain a consistent system load, once a core completes its alignment, it immediately begins a new alignment of the dataset if other cores are still processing their first alignment. This process continues until every core has completed at least one full alignment. Only the execution time for the first alignment on each core is included in the analysis. Multi-threaded speedup is then computed relative to single-threaded BiWFA performance by dividing the single-thread BiWFA execution time (reported in the previous section) by the scaled time observed for each algorithm using *N* threads. BiWFA serves as the baseline for speedup calculations because it consistently completes all test cases without exhausting memory, whereas KSW2, KSW2+Singletrack, WFA, and WFA+Singletrack run out of memory in certain scenarios.

Figure 3 shows the speedup of each algorithm relative to BiWFA single-threaded execution for thread counts of 1, 14, 28, 42, 56, 70, 84, 98, and 112. In this figure, rows correspond to datasets, and columns to cost functions. That is, each subplot corresponds to a dataset–function pair. Note that the y-axis range varies across subplots. The computing node consists of two chips: thread counts up to 56 utilise a single chip, while higher thread counts engage both chips. This architectural transition is indicated by a thick vertical line at 56 threads. In some cases, the speedup curves end before reaching 112 threads because the algorithm exhausts the node’s available memory. Because each core performs the same workload as in the single-threaded benchmarks, total memory usage at a given thread count can be precisely computed by multiplying the per-thread memory reported in Table 2 by the number of threads.

**Figure 3.**
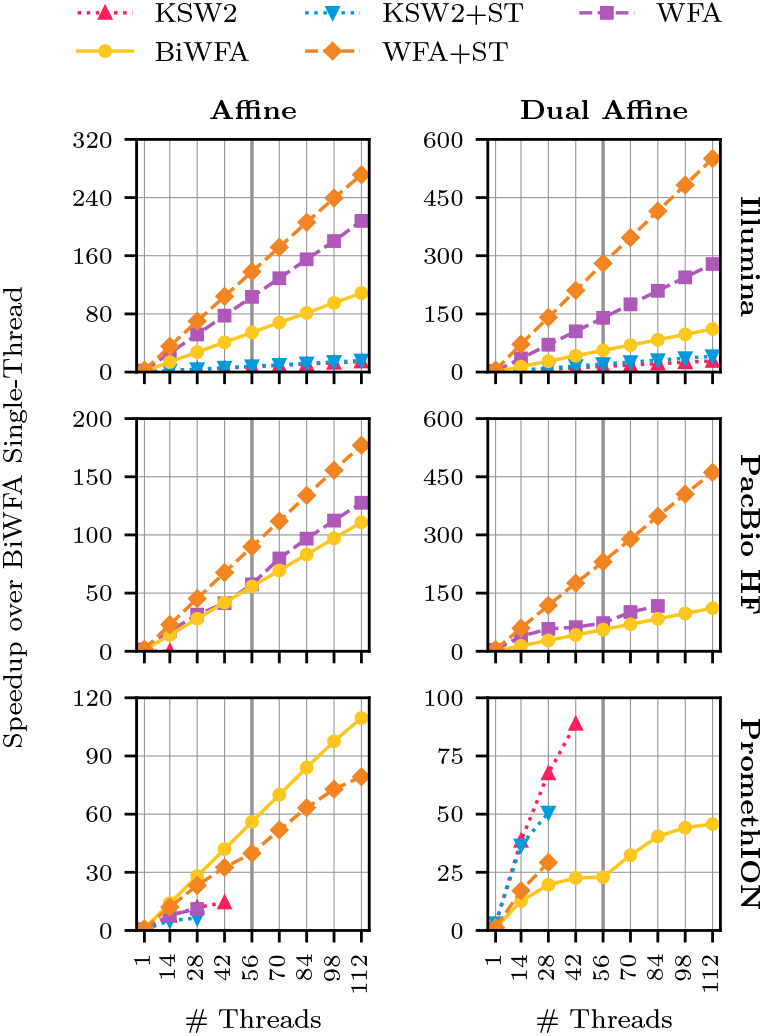
Multi-threaded speedups of KSW2, KSW2+ST, WFA, BiWFA, and WFA+Singletrack relative to single-threaded BiWFA (ST = Single-track).

The parallel performance of each algorithm is influenced by the use of shared resources, such as the last-level cache (LLC) and main memory. When alignment tasks fit entirely within the private L1 and L2 caches of each core, the algorithms achieve ideal speedups: the scaled execution time with *N* threads matches that of single-threaded execution. This is the case for all algorithms on the Illumina dataset, which comprises short sequences.

As memory requirements increase, particularly with the dual gap-affine cost function, WFA, KSW2, and KSW2+Singletrack increasingly rely on shared resources, causing deviations from ideal performance. In contrast, both BiWFA and WFA+Singletrack maintain ideal or near-ideal scaling in most cases due to their more efficient utilization of the memory hierarchy. Consequently, and consistent with the single-threaded performance results discussed previously, WFA+Singletrack outperforms both BiWFA and WFA in the majority of executions in which it does not exhaust the available memory. Notably, WFA+Singletrack’s efficient memory usage enables higher thread counts on large datasets than WFA. For example, when aligning the PacBio HiFi dataset using the dual-gap affine scoring, WFA+Singletrack can utilize all 112 cores on the node, whereas WFA exhausts available memory beyond 84 threads.

KSW2+Singletrack achieves better parallel performance than KSW2 at low thread counts for the dual-gap affine cost function, consistent with the single-threaded results. On the other hand, KSW2 shows the best parallel performance among the studied algorithms across 14–42 threads on the PromethION dataset. While the lower memory requirements of KSW2 allow it to reach higher thread counts than KSW2+Singletrack, the memory demands of both algorithms ultimately limit their scalability, leading to memory exhaustion at lower thread counts than WFA+Singletrack and BiWFA.

At equivalent thread counts, WFA+Singletrack achieves up to a 3.2× speedup over WFA and a maximum speedup of 5.2× over BiWFA. Meanwhile, KSW2+Singletrack reaches speedups of up to 1.4× over KSW2 at low thread counts, while KSW2 exhibits better scalability and is thus preferable for fully loaded nodes.

## 7 Conclusion

In this work, we present Singletrack, a general backtrace algorithm that enables the computation of optimal gap-affine and dual gap-affine alignments with low memory requirements (comparable to gap-linear alignments), without adding alignment overheads, and with a simple, straightforward integration into existing tools.

We implement Singletrack in the KSW2 and WFA2-lib libraries. Our results demonstrate that Singletrack achieves equal memory usage for gap-affine and dual gap-affine alignments in KSW2, and reduces memory consumption by 3× and 5×, respectively, compared to the original WFA implementation. Moreover, Singletrack enables performance improvements during the alignment step, resulting in speedups of up to 2.1× over the baseline versions of KSW2 and WFA2-lib. We also compared our Singletrack-enhanced WFA to BiWFA, which employs a divide-and-conquer approach to achieve linear memory usage. Our experiments show that WFA with Singletrack achieves up to 5.2× higher performance than Bi-WFA, at the cost of increased memory consumption, making it a compelling option for aligning moderately large sequences.

In summary, our method offers a practical solution to overcome the memory limitations of gap-affine and dual gap-affine sequence alignment, while also providing additional performance improvements during alignment. Moreover, Singletrack’s lightweight implementation integrates easily with existing alignment tools, enhancing their practicality and availability for the bioinformatics community. We expect Singletrack to contribute to the development of more efficient and scalable bioinformatics tools that can handle large-scale sequencing datasets in the future.

## 8 Funding

This work was supported by the Spanish Ministry of Science and Innovation MCIN/AEI/10.13039/501100011033 (contracts PID2020-113614RB-C21, PID2022-136454NB-C22, PID2023-146193OB-I00, and PID2023-146511NB-I00), by the Generalitat de Catalunya GenCat-DIUiE (GRR) (grant number 2021-SGR-00763 and 2021-SGR-00574), by the Gobierno de Aragón (E45_20R T58_23R research groups), the Arm-BSC Center of Excellence, by Lenovo-BSC Contract-Framework Contract (2022), the Spanish Ministry, Ministerio para la Transformación Digital y de la Función Pública, and the European Union – NextGenerationEU through the Càtedra Chip UPC project (grant number TSI-069100-2023-0015), the European Union’s Horizon Europe Programme under the STRATUM Project (grant agreement no. 101137416), and by the Barcelona Zettascale Laboratory, backed by the Ministry for Digital Transformation and of Public Services, within the framework of the Recovery, Transformation, and Resilience Plan funded by the European Union – NextGenerationEU. The funders had no role in study design, data collection and analysis, decision to publish, or preparation of the manuscript.

## Footnotes

1 KSW2 available at https://github.com/lh3/ksw2

2 https://github.com/smarco/WFA2-lib

3 https://github.com/genome-in-a-bottle/giab_data_indexes

4 https://precision.fda.gov/challenges/10/intro

## Notes

### Competing Interest Statement

The authors have declared no competing interest.

