## Supplementary Material for "Singletrack: An Algorithm for Improving Memory Consumption and Performance of Gap-Affine Sequence Alignment"

Lorién López-Villellas[1], Cristian Iñiguez[2], Albert Jiménez-Blanco[2], Quim Aguado-Puig[4], Miquel Moretó[3,2], Jesús Alastruey-Benedé[1], Pablo Ibáñez[1], and Santiago Marco-Sola[3,2]

[1]Departamento de Informática e Ingeniería de Sistemas, Universidad de Zaragoza, Spain
[2]Barcelona Supercomputing, Spain
[3]Department of Computer Science, Universitat Politècnica de Catalunya, Spain
[4]Departament d'Arquitectura de Computadors i Sistemes Operatius, Universitat Autònoma de Barcelona, Spain

**Summary**

This supplementary material complements the main manuscript by presenting the extended classical gap-affine equations and the classical backtrace algorithm for gap-affine. It also details the dual gap-affine recurrence relations for the classical formulation, the Suzuki-Kasahara formulation, and the wavefront alignment algorithm (WFA). In addition, it provides a detailed description of the Singletrack backtrace algorithm as applied to the WFA.

### 1 Gap-Affine and Dual Gap-Affine Extended Equations

Equations 1 and 2 present the complete recurrence relations, including the computation of boundary cells, for the DP matrices used in the gap-affine and dual gap-affine scoring functions, respectively.

$$
\begin{aligned}
I1_{i,j} &= \begin{cases} \infty & (i \geq 0, j = 0) \\ o_1 + j \cdot e_1 & (i = 0, j > 0) \\ \min \begin{cases} I1_{i,j-1} + e_1 \\ M_{i,j-1} + o_1 + e_1 \end{cases} & (i > 0, j > 0) \end{cases} \\
D1_{i,j} &= \begin{cases} o_1 + i \cdot e_1 & (i > 0, j = 0) \\ \infty & (i = 0, j \geq 0) \\ \min \begin{cases} D1_{i-1,j} + e_1 \\ M_{i-1,j} + o_1 + e_1 \} \end{cases} & (i > 0, j > 0) \end{cases} \\
M_{i,j} &= \begin{cases} 0 & (i = 0, j = 0) \\ o_1 + i \cdot e_1 & (i > 0, j = 0) \\ o_1 + j \cdot e_1 & (i = 0, j > 0) \\ \min \begin{cases} I1_{i,j} \\ D1_{i,j} \\ M_{i-1,j-1} + S(i,j) \end{cases} & (i > 0, j > 0) \end{cases}
\end{aligned} \tag{1}
$$

$$
\begin{aligned}
I1_{i,j} &= \begin{cases} \infty & (i \geq 0, j = 0) \\ o_1 + j \cdot e_1 & (i = 0, j > 0) \\ \min \begin{cases} I1_{i,j-1} + e_1 \\ M_{i,j-1} + o_1 + e_1 \end{cases} & (i > 0, j > 0) \end{cases} \\
D1_{i,j} &= \begin{cases} o_1 + i \cdot e_1 & (i > 0, j = 0) \\ \infty & (i = 0, j \geq 0) \\ \min \begin{cases} D1_{i-1,j} + e_1 \\ M_{i-1,j} + o_1 + e_1 \end{cases} & (i > 0, j > 0) \end{cases} \\
I2_{i,j} &= \begin{cases} \infty & (i \geq 0, j = 0) \\ o_2 + j \cdot e_2 & (i = 0, j > 0) \\ \min \begin{cases} I2_{i,j-1} + e_2 \\ M_{i,j-1} + o_2 + e_2 \end{cases} & (i > 0, j > 0) \end{cases} \\
D2_{i,j} &= \begin{cases} o_2 + i \cdot e_2 & (i > 0, j = 0) \\ \infty & (i = 0, j \geq 0) \\ \min \begin{cases} D2_{i-1,j} + e_2 \\ M_{i-1,j} + o_2 + e_2 \end{cases} & (i > 0, j > 0) \end{cases} \\
M_{i,j} &= \begin{cases} 0 & (i = 0, j = 0) \\ \min \begin{cases} o_1 + i \cdot e_1 \\ o_2 + i \cdot e_2 \end{cases} & (i > 0, j = 0) \\ \min \begin{cases} o_1 + j \cdot e_1 \\ o_2 + j \cdot e_2 \end{cases} & (i = 0, j > 0) \\ \min \begin{cases} I1_{i,j} \\ D1_{i,j} \\ I2_{i,j} \\ D2_{i,j} \\ M_{i-1,j-1} + S(i,j) \end{cases} & (i > 0, j > 0) \end{cases}
\end{aligned}
\tag{2}
$$

#### 2 Classical Backtrace Algorithm

Algorithm 1 presents the classical backtrace algorithm for the gap-affine model, which requires access to all three DP matrices: $M$, $I1$, and $D1$.

#### 3 Dual Gap-Affine Equations in Suzuki-Kasahara Formulation

Equation 3 shows the recurrence relations for the dual gap-affine scoring function in the Suzuki-Kasahara formulation. Note that the $A$ matrix is not stored; it is only used as a temporary variable to compute $\Delta H$ and $\Delta V$.

**Algorithm 1** Classical gap-affine backtrace

**Input** Sequences $q_{0,\dots,n-1}$, $t_{0,\dots,m-1}$, matrices $M, I1, D1$, penalties $\{a, x, o_1, e_1\}$
**Output** CIGAR string that represents the optimal alignment

```
function BACKTRACE(q, t, M, I, D, p = {a, x, o_1, e_1})
    Definition: PUSH_OP(cigar, op, len) appends op len
    times to cigar.
    cigar ← ϵ, i ← n, j ← m, state ← M
    while i > 0 or j > 0 do
        if state = M then
            if M_{i,j} = I_{i,j} then
                state ← I
            else if M_{i,j} = D_{i,j} then
                state ← D
            else
                if q_{i-1} = t_{j-1} then
                    PUSH_OP(cigar, 'M', 1)
                else
                    PUSH_OP(cigar, 'X', 1)
                end if
                i ← i − 1, j ← j − 1
            end if
        else if state = I and j > 0 then
            if I_{i,j} ≠ I_{i,j-1} + e_1 then
                state ← M
            end if
            PUSH_OP(cigar, 'I', 1)
            j ← j − 1
        else if state = D and i > 0 then
            if D_{i,j} ≠ D_{i-1,j} + e_1 then
                state ← M
            end if
            PUSH_OP(cigar, 'M', 1)
            i ← i − 1
        else
            state ← M
        end if
    end while
    return cigar
end function
```

$$
\begin{aligned}
A_{i,j} &= \min \begin{cases} S(i,j) \\ \Delta E1_{i,j-1} + \Delta V_{i,j-1} + e_1 \\ \Delta F1_{i-1,j} + \Delta H_{i-1,j} + e_1 \\ \Delta E2_{i,j-1} + \Delta V_{i,j-1} + e_2 \\ \Delta F2_{i-1,j} + \Delta H_{i-1,j} + e_2 \end{cases} \\
\Delta H_{i,j} &= A_{i,j} - \Delta V_{i,j-1} \\
\Delta V_{i,j} &= A_{i,j} - \Delta H_{i-1,j} \\
\Delta E1_{i,j} &= \min \begin{cases} o_1 \\ \Delta E1_{i,j-1} - \Delta H_{i,j} + e_1 \end{cases} \\
\Delta F1_{i,j} &= \min \begin{cases} o_1 \\ \Delta F1_{i-1,j} - \Delta V_{i,j} + e_1 \end{cases} \\
\Delta E2_{i,j} &= \min \begin{cases} o_2 \\ \Delta E2_{i,j-1} - \Delta H_{i,j} + e_2 \end{cases} \\
\Delta F2_{i,j} &= \min \begin{cases} o_2 \\ \Delta F2_{i-1,j} - \Delta V_{i,j} + e_2 \end{cases}
\end{aligned} \tag{3}
$$

### 4 Dual Gap-Affine Equations in Wavefront Alignment Algorithm

Equation 4 shows the recurrence relations for the dual gap-affine scoring function in WFA.

$$
\begin{aligned}
\widetilde{I1}_{z,k} &= \max \begin{cases} \widetilde{M}_{z-o_1-e_1,k-1} \\ \widetilde{I1}_{z-e_1,k-1} + 1 \end{cases} \\
\widetilde{D1}_{z,k} &= \max \begin{cases} \widetilde{M}_{z-o_1-e_1,k+1} \\ \widetilde{D1}_{z-e_1,k+1} \end{cases} \\
\widetilde{I2}_{z,k} &= \max \begin{cases} \widetilde{M}_{z-o_2-e_2,k-1} \\ \widetilde{I2}_{z-e_2,k-1} + 1 \end{cases} \\
\widetilde{D2}_{z,k} &= \max \begin{cases} \widetilde{M}_{z-o_2-e_2,k+1} \\ \widetilde{D1}_{z-e_2,k+1} \end{cases} \\
\widetilde{M}_{z,k} &= \max \begin{cases} \widetilde{M}_{z-x,k} + 1 \\ \widetilde{I1}_{z,k} \\ \widetilde{D1}_{z,k} \\ \widetilde{I2}_{z,k} \\ \widetilde{D2}_{z,k} \end{cases}
\end{aligned} \tag{4}
$$

### 5 Extended Description of Singletrack in Wavefront Alignment Algorithm

In the main text, we used the Wavefront Alignment Algorithm [1] (WFA) as a case study to evaluate the impact of the Singletrack backtrace algorithm. Here, we provide specific details for integrating Singletrack into WFA. We focus on the gap-affine scoring model, but the approach generalizes seamlessly to the dual gap-affine score function.

Analogous to the general Singletrack strategy, we assume that the WFA backtrace algorithm only has access to the $\widetilde{M}_{0,\ldots,s}$ wavefronts, where $s$ denotes the optimal alignment score. It is important to note that WFA does not explicitly store all computed DP cells due to its use of the `extend` operation. After generating a DP cell with score $z$, diagonal $k$, and offset $off'$ using the `next` operation, the `extend` operation increases the offset as long as matches are found along the diagonal, ultimately storing only the offset $off$ corresponding to the furthest-reaching cell for each diagonal and score. Consequently, when considering a current cell $off = \widetilde{M}_{z,k}$, the original offset $off'$ before extension is not directly available. To address this, the Singletrack algorithm performs a reverse extension (`rev_extend`) operation. Starting from the current cell $off = \widetilde{M}_{z,k}$, `rev_extend` decrements the offset while matches are found along the diagonal in reverse, recovering the tentative original offset $off'$. Once this reverse extension is complete, we determine the origin of the predecessor cell by checking if $off'$ is consistent with the expected predecessor in the $M$ matrix, specifically whether $off'$ equals $\widetilde{M}_{z-x,k} + 1$ for a substitution. If this condition is satisfied, the predecessor cell is in the $M$ matrix (i.e., in a $\widetilde{M}$ wavefront), and the algorithm continues the backtrace from this position. If the condition is not true, the predecessor must be in one of the indel wavefronts, either $\widetilde{I}_1$ or $\widetilde{D}_1$. At this stage, we open two tentative backtrace paths: one assuming the current cell originated from $\widetilde{I}_1$, and the other from $\widetilde{D}_1$. For these tentative paths, the offset $off'$ obtained from `rev_extend` may not correspond to the original offset of $\widetilde{M}z - o_1 - e_1, k$. The real original offset could be any value between $off'$ and $off$, since the insertion or deletion may have occurred in the middle of a chain of consecutive matches along the diagonal. Knowing this, the algorithm increases the gap length $l$ iteratively, computing the expected score $z'$ and checking for a consistent path back to $M$. In the case of insertions, this is done by verifying if $off'$ is less than or equal to $\widetilde{M}_{z',k-l} + l$ and less than or equal to $off$. For deletions, the check is if $off'$ is less than or equal to $\widetilde{M}_{z',k+l}$ and less than or equal to $off$. When one of these criteria is satisfied, the corresponding path is confirmed as the correct backtrace path, and the other path is discarded. The algorithm then resumes the backtrace from the identified $M$ cell. The pseudocode for the Singletrack backtrace algorithm in WFA is presented in Algorithm 2.

We reuse the example from the main text to demonstrate the Singletrack backtrace in WFA. Figure 1 shows the DP cells and wavefronts computed by WFA for the alignment of sequences $q =$ `GCA` and $t =$ `GCCAA`. The $\widetilde{I}_1$ and $\widetilde{D}_1$ wavefronts are shown to provide full context for the example, although they are never used by the Singletrack backtrace algorithm itself. The alignment begins at cell $M_{3,5}$ (the current cell), which corresponds to $\widetilde{M}_{10,2} = 5$. Here, $z = s = 10$, and the diagonal and offset are $k = 2$ and $off = 5$, respectively. Since we start in $M$, the Singletrack algorithm performs the `rev_extend` operation. In this case, there is one match (the letter `A`) on diagonal $k = 2$ starting at $off = 5$, so $off' = 4$. However, the cell preceding $\widetilde{M}_{10,2}$ before executing the `extend` operation was not generated from a substitution, since $off' \neq \widetilde{M}_{z',2}+1$, with $z' = z-x = 6$.

**Algorithm 2** Gap-affine Singletrack backtrace in WFA

**Input** Sequences $q_{0,\ldots,n-1}$, $t_{0,\ldots,m-1}$, optimal score $s$, list of wavefronts $\widetilde{M}_{0,\ldots,s}$, penalties $\{a = 0, x > 0, o_1 > 0, e_1 > 0\}$

**Output** CIGAR string that represents the optimal alignment

**function** $\textsc{Backtrace}(q, t, s, \widetilde{M}_{0,\ldots,s}, p = \{a = 0, x > 0, o_1 > 0, e_1 > 0\})$
    **Definition:** `PUSH_OP(cigar, op, p)` adds `op` to `cigar` `p` times.
    **Definition:** $S(i, j) = a$ if $q_{i-1} = t_{j-1}$, $x$ otherwise.
    ▷ For simplicity, we assume that an invalid access to $\widetilde{M}_{0,\ldots,s}$ (either invalid score or diagonal) returns $-\infty$
    $\text{cigar} \leftarrow \epsilon, z \leftarrow s, l \leftarrow 0, k \leftarrow m - n, off \leftarrow m, off' \leftarrow 0, \text{state} \leftarrow M$
    **while** $s > 0$ **do**
        **if** $\text{state} = M$ **then**
            $i \leftarrow off - k, j \leftarrow off, off' \leftarrow off$
            **while** $i > 0$ **and** $j > 0$ **and** $q_{i-1} = t_{j-1}$ **do** ▷ Reverse extend()
                $i \leftarrow i - 1, j \leftarrow j - 1, off' \leftarrow off' - 1$
            **end while**
            $z' \leftarrow z - x$
            **if** $\widetilde{M}_{z',k} + 1 = off'$ **then**
                $\textsc{push\_op}$(cigar, ‘M’, $off' - off$)
                $\textsc{push\_op}$(cigar, ‘X’, 1)
                $z \leftarrow z', off = off'$
            **else**
                $l \leftarrow 0, \text{state} \leftarrow \bar{M}$
            **end if**
        **else**
            $l \leftarrow l + 1, z' \leftarrow z - o_1 - l \cdot e_1$
            **if** $off' \leq \widetilde{M}_{z',k-l} + l \leq off$ **then**
                $\textsc{push\_op}$(cigar, ‘M’, $off - (\widetilde{M}_{z',k-l} + l)$)
                $\textsc{push\_op}$(cigar, ‘I’, $l$)
                $off \leftarrow \widetilde{M}_{z',k-l}, k \leftarrow k - l, z \leftarrow z', \text{state} \leftarrow M$
            **else if** $off' \leq \widetilde{M}_{z',k+l}\widetilde{M}_{z',k+l} \leq off$ **then**
                $\textsc{push\_op}$(cigar, ‘M’, $off - \widetilde{M}_{z',k+l}$)
                $\textsc{push\_op}$(cigar, ‘I’, $l$)
                $off \leftarrow \widetilde{M}_{z',k+l}, k \leftarrow k + l, z \leftarrow z', \text{state} \leftarrow M$
            **end if**
        **end if**
    **end while**
    $\textsc{push\_op}$(cigar, ‘M’, $off$)
    **return** cigar
**end function**

Therefore, the predecessor cell must be in either $I1$ or $D1$. At this point, the gap length $l$ is initialized to 0 and then increased to 1, and two tentative paths have been opened. The following conditions are then checked: $off' \leq \widetilde{M}_{z',k-l} + l \leq off$ for the insertion path, and $off' \leq \widetilde{M}_{z',k+l} \leq off$ for the deletion path, where $z' = z - o_1 - l \cdot e_1 = 10 - 6 - 1 \cdot 2 = 2$. None of the conditions is true, so the gap length $l$ is increased to 2. In this case, the condition $off' \leq \widetilde{M}_{z',k-l} + l \leq off$, where $z' = z - o_1 - l \cdot e_1 = 10 - 6 - 2 \cdot 2 = 0$ is satisfied, indicating that the optimal path to $M_{3,5}$ goes through $M_{2,2}$, $I1_{2,3}$, and $M_{2,4}$. At this stage, a match and two insertions are added to the CIGAR string, which is initialized as `IIM`. Upon reaching cell $M_{2,2}$ with score $z = 0$, the condition of the `while` loop is no longer satisfied, and the loop exits. Since $off > 0$, there remain $off$ matches to be added to the CIGAR string, which are appended before completing the backtrace process. The final CIGAR string is thus `MMIIM`.

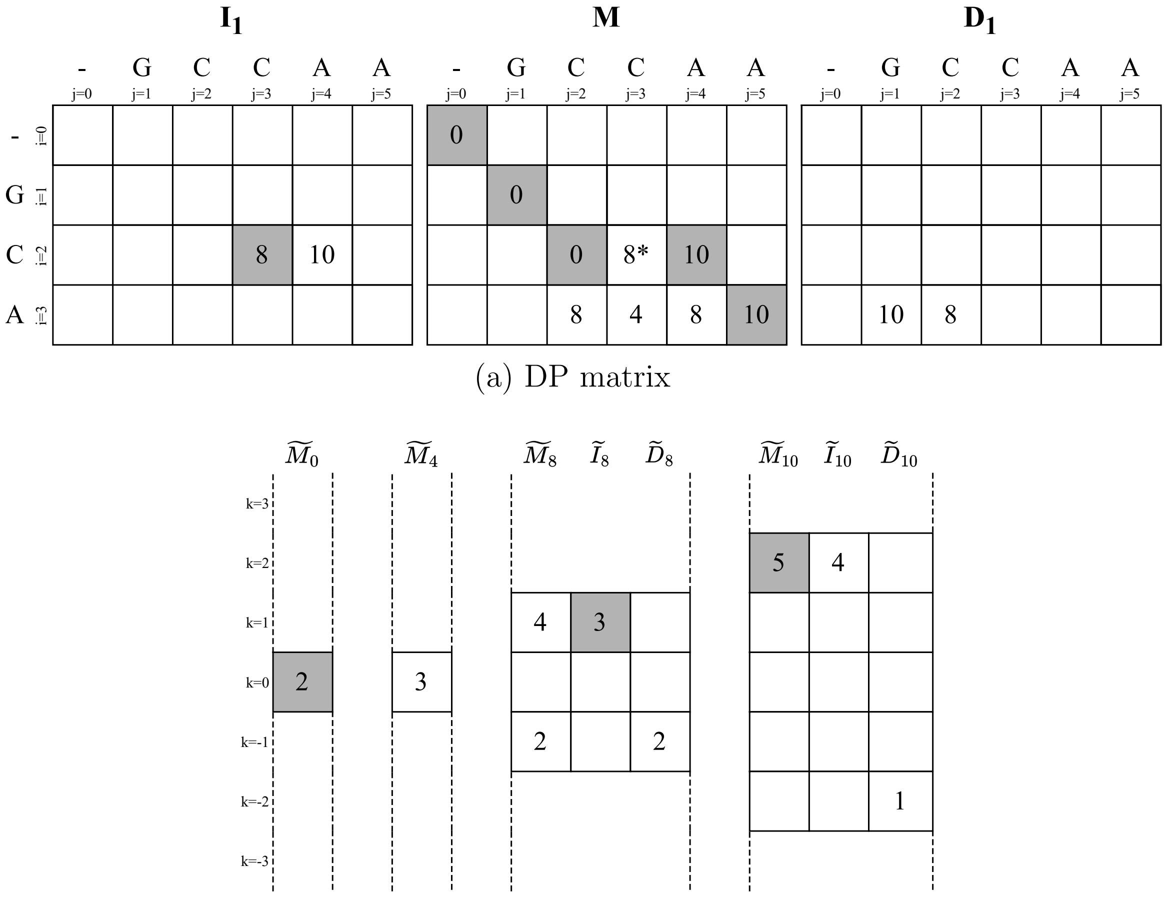

(a) DP matrix

(b) Wavefronts

Figure 1: Computed cells of the DP matrix (a) and wavefronts (b) generated by WFA during the alignment of sequences $q$ = `GCA` and $t$ = `GCCAA` using the gap-affine cost function with penalties $a = 0$, $x = 4$, $o_1 = 6$, and $e_1 = 2$. DP matrix cells marked with an asterisk (*) indicate cells that are computed but not stored due to the `extend` operation. Empty cells in the wavefronts represent non-existent DP cells; that is, for a given score $z$ and diagonal $k$, no DP cell exists with that score on that diagonal. Note that only a subset of the $\widetilde{I}_1$ and $\widetilde{D}_1$ wavefronts needs to be stored during alignment, as the Singletrack backtrace algorithm requires only the $\widetilde{M}$ wavefronts. Dark-shaded cells indicate matrix entries that are part of the optimal alignment path.
